# Demonstration of Patient-Specific Simulations To Assess Left Atrial Appendage Thrombogenesis Risk

**DOI:** 10.1101/2020.05.07.083220

**Authors:** Manuel García-Villalba, Lorenzo Rossini, Alejandro Gonzalo, Davis Vigneault, Pablo Martinez-Legazpi, Oscar Flores, Javier Bermejo, Elliot McVeigh, Andrew M. Kahn, Juan C. del Álamo

## Abstract

Atrial fibrillation (AF) alters left atrial (LA) hemodynamics, which can lead to thrombosis in the left atrial appendage (LAA), systemic embolism and stroke. A personalized risk-stratification of AF patients for stroke would permit improved balancing of preventive anticoagulation therapies against bleeding risk. We investigated how LA anatomy and function impact LA and LAA hemodynamics, and explored whether patient-specific analysis by computational fluid dynamics (CFD) can predict the risk of LAA thrombosis. We analyzed 4D-CT acquisitions of LA wall motion with an in-house immersed-boundary CFD solver. We considered six patients with diverse atrial function, three without a LAA thrombus (LAAT/TIA-neg), and three with either a LAA thrombus (removed digitally before running the simulations) or a history of transient ischemic attacks (LAAT/TIA-pos). We found that blood inside the left atrial appendage of LAAT/TIA-pos patients had marked alterations in residence time and kinetic energy when compared with LAAT/TIA-neg patients. In addition, we showed how the LA conduit, reservoir and booster functions distinctly affect LA and LAA hemodynamics. While the flow dynamics of fixed-wall and moving-wall simulations differ significantly, fixed-wall simulations risk-stratified our small cohort for LAA thrombosis only slightly worse than moving-wall simulations.

## 1. Introduction

Atrial fibrillation (AF) is the most common cardiac arrhythmia, affecting approximately 35 million people worldwide [1]. During AF, the atria beat irregularly and their function is not coordinated with the ventricles. Each atrium has a morphologically characteristic body, and an appendage where thrombi form preferentially [10]. Because some of these thrombi can travel to the brain, the risk of stroke of patients with AF is five times higher compared to the general population, and AF causes 15% of all strokes. Anticoagulation drugs reduce the risk of strokes in patients with AF. However, because these drugs are associated with an increased risk of bleeding, they are prescribed only to patients for whom the risk of stroke outweighs the bleeding risk. Current methods to risk-stratify patients are not personalized and contain no information about the patients’ cardiac anatomy or blood flow; instead, they are based on demographic and clinical factors such as age, gender, or coexistent hypertension. These factors are derived from large clinical trials and have limited predictive value for a specific patient. Furthermore, there are currently many patients for whom there is clinical uncertainty as to whether anticoagulation is beneficial [13].

Because blood stasis is considered necessary for thrombogenesis, we focus on the fluid mechanics of the left atrium (LA) and the left atrial appendage (LAA). The LA is a compliant structure that performs several functions during the cardiac cycle. It operates as a reservoir for the incoming flow through the pulmonary veins (PVs), as a conduit during early left ventricle (LV) diastole, and as a booster pump that augments LV filling during late LV diastole [12]. As a result of the LA complex geometry and motion, and its variability among the population, the flow dynamics within this chamber are complex, and thus far have not been fully characterized. Compared to the LV, there are few patient-specific computational fluid dynamics (CFD) analyses of the LA. In fact, early CFD studies of LA hemodynamics were part of whole-left-heart simulations focusing mostly on how LA geometry affects the LV flow features [28].

CFD analyses focusing on the LA are relatively recent. Koizumi et al [15] prescribed synthetic wall motions of the LA of a healthy subject to study how atrial function may contribute to washing out the LAA. Several other works followed a similar approach, offering somewhat contradictory results. Masci et al [18] studied two subjects with a history of AF, comparing the effects of LA wall motion in sinus rhythm with weak random motion, and concluding that reduced LAA contractility can result in slow blood flow. By contrast, Otani et al [22] studied two subjects with impaired atrial function and a history of AF but no evidence of prior stroke or TIAs, showing that the subject with better atrial function experienced more LAA blood stasis despite having higher flow velocities.

It appears that LAA morphology correlates with the risk of stroke in patients with AF [2, 5, 29]. However, the degree of correlation is only moderate for the most commonly observed LAA shapes and the mechanisms responsible for this correlation are not well understood. Several CFD studies have explored the effect of LAA geometry on blood flow but, taken together, have offered ambiguous results. Bosi et al [3] found that LAA morphologies that are clinically associated with higher stroke incidence have impaired blood washout, whereas other studies [8, 19] reported that LAA stasis depends critically on additional factors such as PV inflow patterns.

Although the ambiguity in previous CFD analyses of LA hemodynamics could be attributed to their reduced sample sizes, it also suggests that blood transit in the LAA is a complex multi-factorial process that cannot be parameterized easily. Considering these challenges, previous efforts have been limited by their lack of a direct validation endpoint to test whether blood stasis increases the risk of LAA thrombosis. In this study, we performed patient-specific CFD simulations in a group of patients with varying atrial function and LAA geometries. Albeit still small (N = 6), this group is larger than those typically considered in CFD studies so far. Importantly, a subgroup of patients (N=3) had either mural LAA thrombosis, which was segmented out prior to the simulations, or a history of transient ischemic attacks (TIAs). This approach offered direct insight into the relationship between atrial flow and LAA thrombosis. Moreover, we considered both moving-wall and fixed-wall anatomies to evaluate the dependence of blood stasis predictions on LA wall motion.

## 2. Materials and Methods

### 2.1. Imaging of Human Subjects

We retrospectively studied N = 6 patients. All of them were enrolled from a database of patients with existing computed tomography (CT) scans with full RR coverage over the entire LA and LAA. Selection of patients was random with the constraint of covering a wide range of LA ejection fractions as assessed by CT. Cardiac-gated cine CT scans were acquired at the National Institutes of Health (NIH), Bethesda, Maryland (N = 3) and at the University of California San Diego (UCSD), California (N = 3). The studies were approved by the Institutional Review Board at both centers.

Imaging was performed following standard clinical protocols at each participating center. Specifically, electrocardiogram (ECG) gating with inspiratory breath-hold was used to sample one entire heartbeat. The images were acquired with 3 different scanner models: 2 studies were obtained on a Siemens Force system (at NIH), 3 studies were obtained on the Toshiba Aquillion ONE (1 and NIH, 2 at UCSD) and one study was obtained on the 256-slice GE Healthcare Revolution CT at UCSD. For each image iodinated contrast was injected followed by saline flush. The doses and injection rates of contrast were chosen based on the patient’s weight using standard clinical protocols.

The images were reconstructed using the manufacturers’ standard algorithms (Toshiba reconstruction filter AIDR3D, FC08-cfa, GE Healthcare AsirV=0.5 standard kernel, Siemens filter kernel br36d). This procedure yielded DICOM files containing z-stacks of images of 512 x 512 pixels with in-plane pixels dimensions of between 0.32 mm and 0.48 mm in the x-y axial plane. Each z-stack contained between 100 to 320 slices with resolutions of 0.5 mm to 1 mm in the z-direction. The multi-phase reconstructions were performed at regular time intervals across the heart cycle, with 5% RR and 10% RR being the minimum and maximum intervals.

### 2.2. Image Processing

The 4D CT scans described above were imported into itk-SNAP [30] and a segmentation was created using the semi-automatic active contour segmentation method. The PVs were clipped by manually choosing planes for each time frame; similarly, the LV and aorta were clipped off and the LAA was identified. From each 3D segmentation, a surface triangulation was extracted in MATLAB, smoothed and resampled to the resolution required for the CFD using iso2mesh [23]. For each subject, the point cloud of vertices of the largest-LA-volume time frame was used as a reference surface and registered to the point clouds from the other time frames using the Coherent Point Drift algorithm [21]. The location of the centroid of each triangle was established by Fourier interpolation with N_FOU_=6 modes to match the temporal resolution of our CFD solver.

Indices of atrial function were calculated from CT images using standard definitions based on chamber volume, as well as on the waveforms of PV and transmitral flow [12] (Table 1). LA sphericity was calculated as described by Bisbal *et al* [2]. The morphology of the LAA was categorized as one of the four types previously proposed by Di Biase *et al* [5] (chicken wing, cactus, windsock or cauliflower) by a cardiologist with expertise in cardiac CT interpretation. The LAA segmentations were skeletonized using the MATLAB image processing toolbox at each reconstructed phase of the cardiac cycle, and the branches of the skeleton were pruned, segmented and labeled automatically (Figure 1). The number of LAA lobes and the length of each lobe were measured (Table 2).

**Table 1.** Summary of anatomical and functional parameters of the left atrium (LA) and left ventricle (LV) in the LAAT/TIA-neg and (left three columns) and LAAT/TIA-pos (right three columns) groups. EF: ejection fraction; PVs: blood volume that enters the LA during LV systole; PVd: blood volume that enters the LA during LV diastole; PVa: blood volume that exits the LA due to reverse flow volume through the pulmonary veins during atrial contraction; E/A ratio: ratio of peak mitral velocities during early diastole (E-wave) and atrial contraction (A-wave).

| Subject Number |  | 1 | 2 | 3 | 4 | 5 | 6 |
| --- | --- | --- | --- | --- | --- | --- | --- |
| LAA Thrombus or history of TIAs |  | No | No | No | Yes (TIAs) | Yes | Yes |
| Sinus Rhythm (at the time of imaging) |  | Yes | Yes | Yes | No | No | No |
| LV Volume (ml) | End-diastolic ($V_{LV,max}$) | 108 | 109 | 147 | 110 | 181 | 208 |
| | End-systolic ($V_{LV,min}$) | 36 | 24 | 49 | 43 | 105 | 77 |
| | Stroke Volume ($SV = V_{LV,max} - V_{LV,min}$) | 72 | 85 | 98 | 67 | 76 | 131 |
| LV EF [$(V_{LV,max} - V_{LV,min}) / V_{LV,max}$] | | 0.67 | 0.78 | 0.67 | 0.61 | 0.42 | 0.63 |
| LA Volume (ml) | Mean | 86.6 | 70.1 | 115 | 145 | 157 | 180 |
| | Minimum ($V_{LA,min}$) | 59.6 | 49.0 | 87.2 | 119 | 150 | 157 |
| | Maximum ($V_{LA,max}$) | 108 | 91.2 | 145 | 155 | 165 | 205 |
| | Pre-A-wave ($V_{LA,pre-A}$) | 94.3 | 72.3 | 119 | 149 | 153 | 178 |
| LA Sphericity [2] |  | 0.80 | 0.82 | 0.78 | 0.86 | 0.83 | 0.81 |
| LA Function assessed by LA volume [12] | Global (LA EF) [$(V_{LA,max} - V_{LA,min}) / V_{LA,max}$] | 0.45 | 0.46 | 0.40 | 0.23 | 0.091 | 0.23 |
| | Expansion Index [$(V_{LA,max} - V_{LA,min}) / V_{LA,min}$] | 0.81 | 0.86 | 0.66 | 0.30 | 0.10 | 0.31 |
| | Passive EF [$(V_{LA,max} - V_{LA,pre-A}) / V_{LA,max}$] | 0.13 | 0.21 | 0.18 | 0.039 | 0.073 | 0.13 |
| | Booster EF [$(V_{LA,pre-A} - V_{LA,min}) / V_{LA,pre-A}$] | 0.37 | 0.32 | 0.27 | 0.20 | 0.020 | 0.12 |
| | Conduit Volume / LV Stroke Volume [$1 - (V_{LA,max} - V_{LA,min}) / SV$] | 0.33 | 0.50 | 0.41 | 0.46 | 0.80 | 0.63 |
| LA Function assessed by PV and transmitral flow [12] | PVs (ml) | 43 | 38 | 51 | 36 | 17 | 49 |
|  | PVd (ml) | 65 | 58 | 60 | 52 | 63 | 94 |
|  | PVs-PVd ratio | 0.66 | 0.66 | 0.84 | 0.69 | 0.27 | 0.52 |
|  | PVa (ml) | 17 | 7.3 | 12 | 13 | 0.14 | 1.4 |
|  | PVa duration (cycles) | 0.108 | 0.106 | 0.133 | 0.117 | 0.039 | 0.068 |
|  | A-wave duration (cycles) | 0.218 | 0.254 | 0.299 | 0.182 | 0.213 | 0.23 |
|  | PVa – A wave duration ratio | 0.49 | 0.43 | 0.44 | 0.64 | 0.18 | 0.30 |
|  | E/A ratio | 1.64 | 2.36 | 3.03 | 2.05 | 3.86 | 3.65 |

**Table 2.**
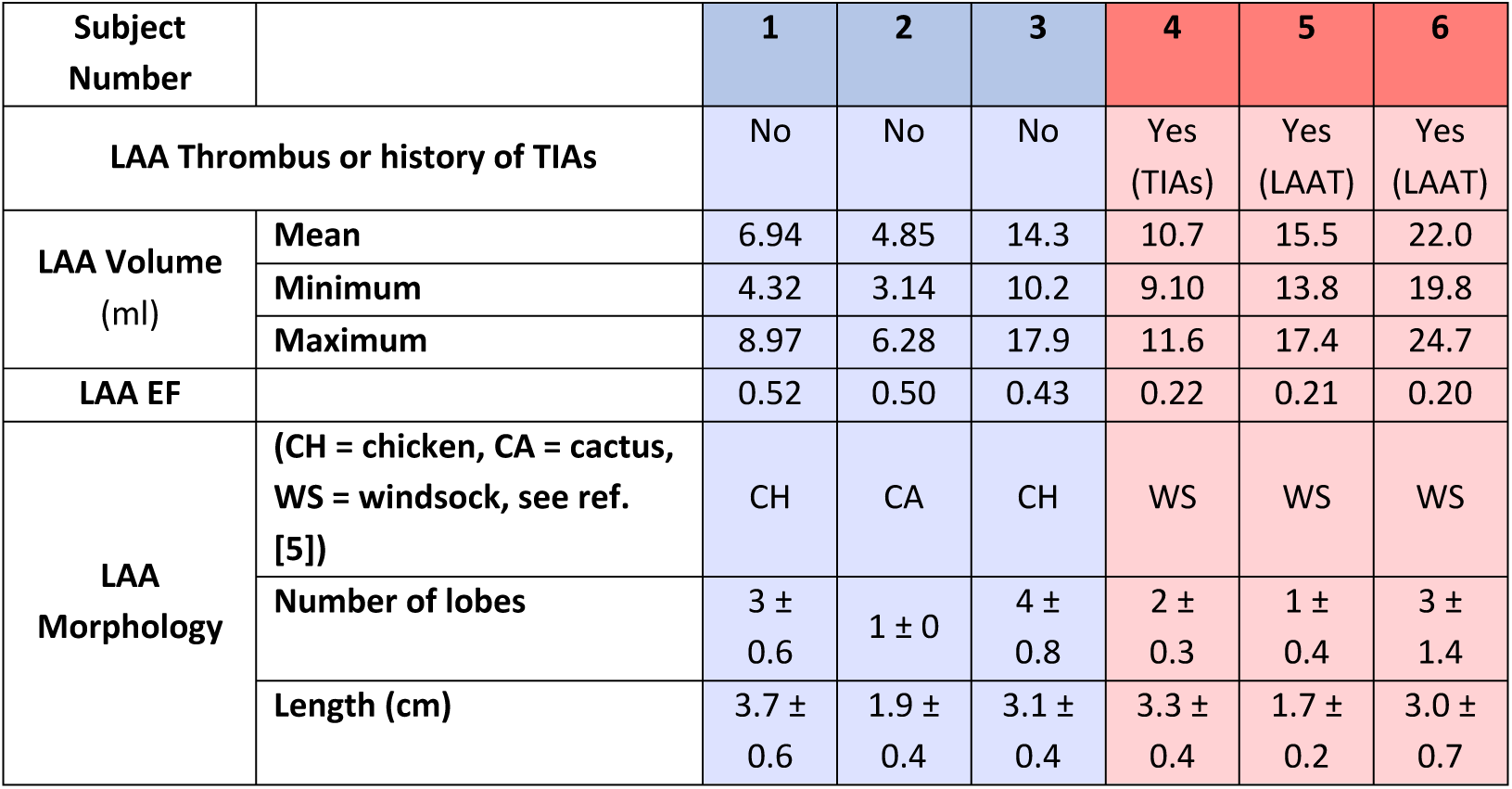
Summary of morphological parameters of the left atrial appendage (LAA) in the LAAT/TIA-neg and (left three columns) and LAAT/TIA-pos (right three columns) groups. Abbreviations: EF: ejection fraction.

**Figure 1.**
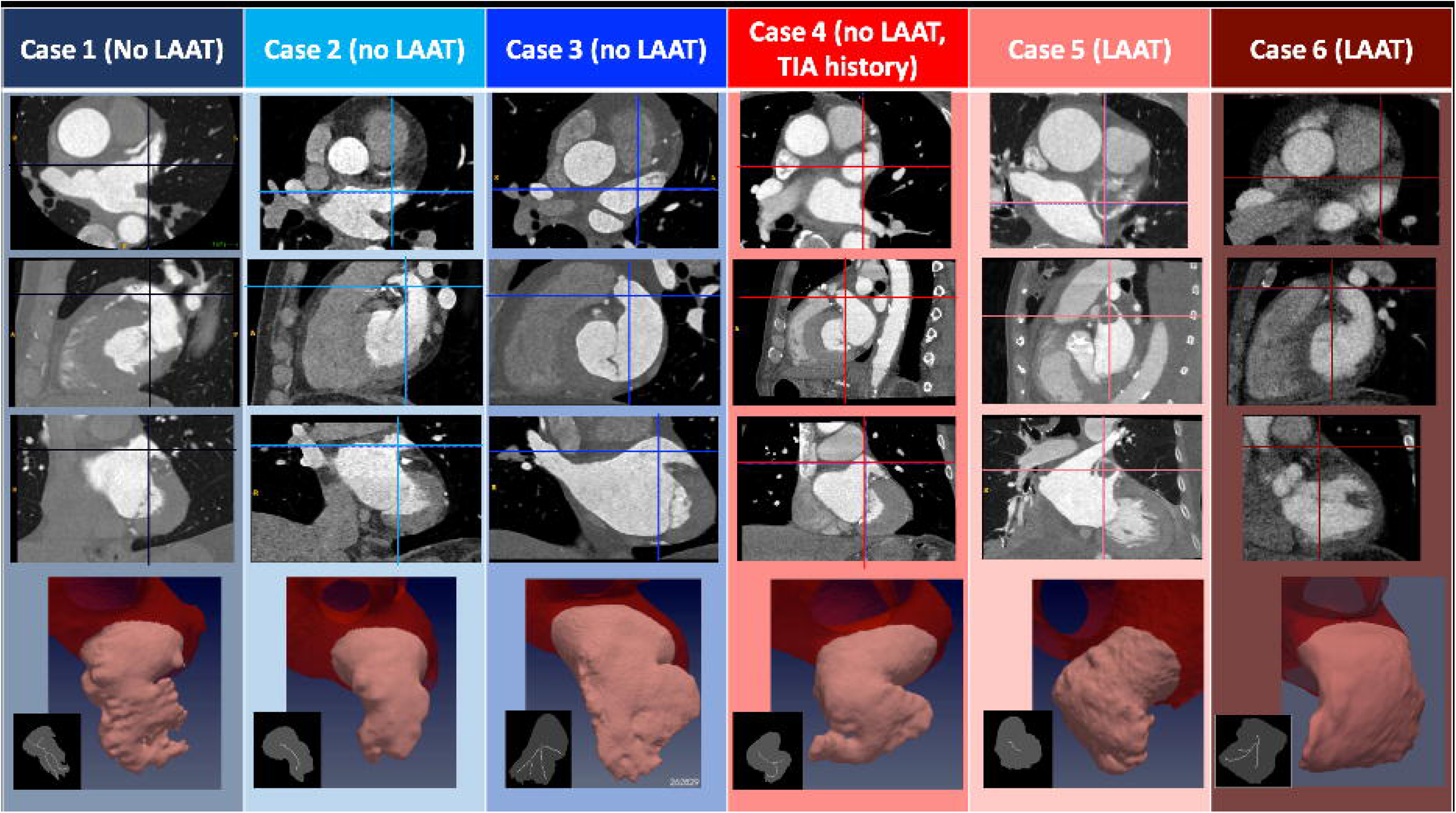
Anatomical Features of Patient-Specific Subjects. The three first rows display CT images corresponding to the N = 6 subjects studied. The images represent, from top to bottom, axial, sagittal and coronal plane sections showing the left atrium, and including LAA thrombi if present (patients 5 and 6). In each panel, the intersections with the other two view planes are represented with a vertical and a horizontal line. The bottom row displays a 3D rendering of the segmented LAA for each subject, and an inset showing its plane projection (gray) and the branches that define the LAA lobes (white). Data are shown at an instant corresponding to 50% of the R-R interval.

### 2.3. Computational Fluid Dynamics

The simulations were performed with the in-house code TUCAN [20], which solves the incompressible Navier-Stokes equations using a fractional step method. The discretization used a staggered Cartesian grid and centered, second-order, finite differences. Time integration was performed with a low-storage, three-stage, semi-implicit Runge–Kutta scheme. Each simulation was initialized from no flow conditions and run for 10 heartbeats at a low resolution (grid spacing Δx=0.090 cm in all directions), with a constant time step Δt and a Courant number *CFL*<0.3. The flow field was then interpolated into a finer grid (Δx=0.051 cm, 256^3^ grid points) and run for another 10 heartbeats with the same *CFL* limit. This resolution is comparable to the finest of the resolutions employed in previous works [28]. In preliminary runs, an even finer grid was employed (Δx = 0.034 cm), showing no relevant differences in the results.

The blood residence time (*T*_*R*_) was computed by solving the forced passive scalar equation [24],

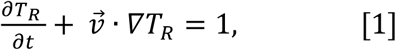

where *T*_*R*_ was set equal to zero at the start of the simulation and at the PVs. Equation 1 was discretized like the Navier-Stokes equations, but spatial fluxes were evaluated with a 3^rd^ order WENO scheme [27]. This scheme ensures a robust solution of the residence time, avoiding Gibbs phenomena near steep gradients without sacrificing accuracy elsewhere.

Wall motion was modelled using the immersed boundary method with the spatio-temporal mesh obtained from the CT images. To specify the inflow boundary conditions through the PVs, we calculated the combined flow rate through all the PVs, *Q*_*PV*_, from mass conservation inside the left atrium. Then, *Q*_*PV*_ was split evenly among all the PVs, resulting in a flow rate through each PV equal to *Q*_*i*_ *= Q*_*PV*_(*t*)/4. We performed preliminary simulations comparing this approach to splitting *Q*_*PV*_ so that the bulk mean velocity was the same in all PVs, and did not find significant differences between the two approaches. The velocity was prescribed at the PVs by applying a volumetric force ***f***_***PV***_ in a buffer region upstream of the PV inlet plane. The target inlet velocity at each PV inlet plane was defined as 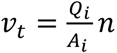, where *A*_*i*_ and ***n*** are the cross-sectional area and the vector normal to each inlet’s plane, respectively. The force was set proportional to the difference between the local velocity and the target inlet velocity, i.e., 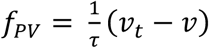, where the constant *τ* is the characteristic time of the forcing. For all simulations presented here, we chose *τ* = 10Δt = 0.5ms, resulting deviations from the target velocity < 5% at the inlet. An analogous scheme was used to set the residence time to zero at the inlet of the PVs. To explore the influence of the motion of the LA wall, we performed corresponding simulations keeping the position of the LA walls fixed at the zero mode of its Fourier temporal series. For the fixed wall-simulations, the A-wave was digitally removed from the transmitral flow, setting *Q*_*MV*_ = 0 for the duration of the atrial contraction.

## 3. Results

### 3.1. Subject population

The median age of the study subjects was 65 (range 50 – 92), and two were female. Subjects 1— 3 were imaged in sinus rhythm and did not have LAA thrombus. Subjects 5 and 6 were imaged in AF with an LAA thrombus, which was digitally removed prior to segmenting the LA geometry for the CFD simulations. Subject 4 was imaged in atrial fibrillation without LAA thrombus, but had a previous history of transient brain ischemic attacks (TIAs). AF was determined based on ECG rhythm strips and lack of late diastolic LA/LAA ejection (“atrial kick”). Based on this information, we categorized subjects 1 – 3 as LAA-thrombus/TIA negative (LAAT/TIA-neg), and subjects 4 – 6 as LAAT/TIA-pos.

Subjects 1 – 3 had normal LV and LA functions as inferred from CT-derived chamber volumes and PV flows (Table 1). Subjects 4 – 6 had enlarged LA with impaired global function. Subject 4 had decreased reservoir function but relatively normal booster function. Subjects 5 and 6 had impaired reservoir and booster function. The LA chamber of subject 5 had severely impaired function, operating mostly as a conduit between the PVs and the LV.

Figure 1 displays primary cardiac CT images of the 6 subjects including segmentations of their LAAs and a plane projection of the morphological skeletonization of the appendage. These images show that subject 5 had an obvious isolated LAA thrombus in the central cavity of the LAA, while subject 6 had the distal section of the LAA filled with thrombus (which was confirmed on a “late” single acquisition obtained 22 seconds after the primary dynamic acquisition). The LAA skeletonization was used to determine LAA lobe number (i.e., the number of skeleton branches) and LAA length (i.e., the length of the branch whose end is most distal to the LAA orifice). Table 2 contains detailed morphological information of the LAA for our study subjects. While the small size of the cohort precludes detailed statistical examination, it is worth noting that the LAAT/TIA-pos subjects had more complex LAA shapes (i.e., windsock vs chicken or cactus) and larger LAA volumes than the LAAT/TIA-neg subjects. We did not observe striking differences in LAA lobe count or LAA length (Table 2).

### 3.2. Flow Visualization

We first present 3D velocity vector maps for one of the subjects with normal atrial function (subject 3, Figure 2). The vectors are colored according to their proximity to four anatomical landmarks: the left PVs, the right PVs, the LAA and the mitral annulus. Three representative instants of the cardiac cycle are shown: LA diastole, LV early filling, and LA systole (Figure 2, top, center and bottom row, respectively). We juxtaposed moving-wall and fixed-wall simulations (Figure 2, left and right column respectively) to illustrate how wall motion affects LA flow dynamics. The flow rate profiles in the PVs and mitral annulus, and the volume profiles of the LA and LAA are also plotted.

**Figure 2.**
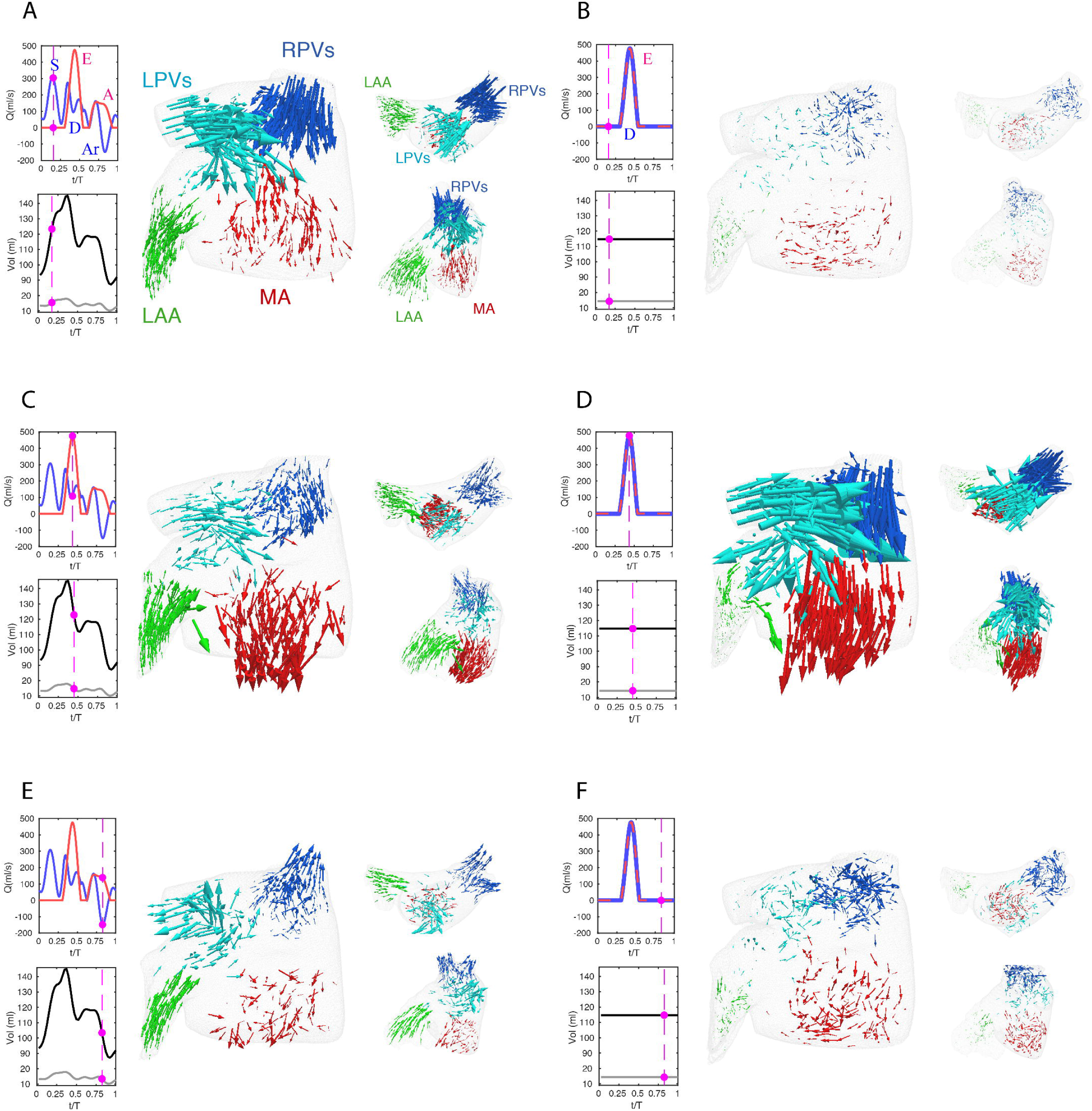
Flow visualization in a left atrium with normally moving and fixed walls. **A, C, D)** Simulation with prescribed patient-specific LA wall motion as obtained from time-resolved CT imaging. **B, D, E)** Simulation in which the position of the LA walls of the same patient were at its time-averaged position. Each panel displays three views of an instantaneous 3-D vector map of blood flow velocity in the left atrium. Clockwise: larger frontal view, smaller axial view and smaller oblique view emphasizing the flow in the left atrial appendage. The vectors are colored according to their proximity to the right pulmonary veins (**blue**), left pulmonary veins (**cyan**), left atrial appendage (**green**) and mitral valve (**red**), and are scaled with the magnitude of blood speed. The panels also include time histories of the flow rate through the mitral valve (**red**) and the cumulative flow rate through the pulmonary veins **(blue)**, and of the volumes of the left atrium (**black**) and the left atrial appendage (**grey**). The magenta bullets indicate the instant of time represented in the vector plots of each panel. The main waves of these profiles are indicated in the plots, i.e., the E-wave and A-wave of transmitral flow during early and late LV filling, the S-wave and D-wave of forward PV flow during LV systole and diastole, and the Ar-wave of backward PV flow during LV diastole, respectively. In the fixed-wall simulations, LA volume is constant and therefore the only transmitral and PV flow waves are the E-wave and the D-wave, and these two waves coincide. The flow vectors inside the LA are represented at three instants of time. **A, B)** *t = 0.16 T*, which corresponds to atrial diastole and peak flow rate through the pulmonary veins in the moving-walls subject. **C, D)** *t = 0.44 T*, which corresponds with left ventricular diastole and peak flow rate through the mitral valve. **E, F)** *t = 0.94 T*, which corresponds with atrial systole and peak backflow rate through the pulmonary vein.

Starting at atrial diastole, the mitral valve is closed, and LA volume is increasing as blood enters the chamber in the moving-wall simulations (Figure 2A, blue arrows). The collision of the left and right PV jets redirects blood towards the mitral annulus (red arrows) and the LAA (green arrows). In contrast, the fixed-wall simulation shows almost no fluid motion at the same timepoint (Figure 2B). Given that the chamber volume does not vary in fixed-wall simulations and the mitral valve is closed, the only possible fluid motion is the remnant flow from ventricular diastole.

During LV relaxation (E-wave, Figure 2C), blood still enters the LA through the PVs forming a jet collision pattern, but LV suction accelerates the blood towards the mitral annulus. Consequently, the flow rate exiting through the mitral annulus (*Q*_*MV*_) exceeds the incoming flow rate through the PVs (*Q*_*PVs*_) and LA volume decreases. The LAA also contracts during this phase, driving a relatively uniform outward flow in its interior. The fixed-wall simulation (Figure 2D) exhibits stronger flow during LV filling even if it was run with the same *Q*_*MV*_. In the moving-wall simulation, passive LA contraction leads to a net difference between the outgoing and incoming flow rates *Q*_*MV*_ - *Q*_*PV*_ ≈ 400 ml/s. However, this difference must be zero in the fixed-wall simulation – thus, PV inflow jets are stronger. The LAA flow pattern in the fixed-wall simulation is reminiscent of a driven cavity flow, consisting of a weak recirculating pattern driven by the flow in the atrial body. This pattern is notably different to the one observed in the moving-wall simulations.

During atrial systole, chamber contraction drives flow into the ventricle (A-wave, Figure 2E) and simultaneously creates backflow into the PVs (Ar-wave). LAA contraction drives a relatively uniform outward flow inside the appendage, similar to the E-wave albeit somewhat slower. In contrast, the fixed-wall simulation displays a weak recirculatory flow corresponding to the late decay stage of the PV and mitral annulus jets generated during LV filling. In particular, the velocities in the LAA are much smaller than in the moving-wall simulation.

### 3.3. The effect of atrial function in flow patterns

The results presented so far reveal that blood flow in the LA and LAA is sensitive to atrial wall motion. However, wall motion can be reduced in different ways depending on whether a patient has impaired reservoir and/or booster function [12]. Thus, we visualized LA and LAA flow patterns across the phases of the cardiac cycle for three patients: one with normal atrial function (subject 3, Figure 3A), one with impaired reservoir function (subject 4, Figure 3B), and one with both impaired reservoir and booster functions (subject 5, Figure 3C). Each snapshot shows blood velocity vectors inside the LAA and iso-surfaces of the second invariant of 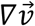, which reveal the vortex systems of the atrial body. Similar to Figure 2, we juxtapose data from moving-wall and fixed-wall simulations.

**Figure 3.**
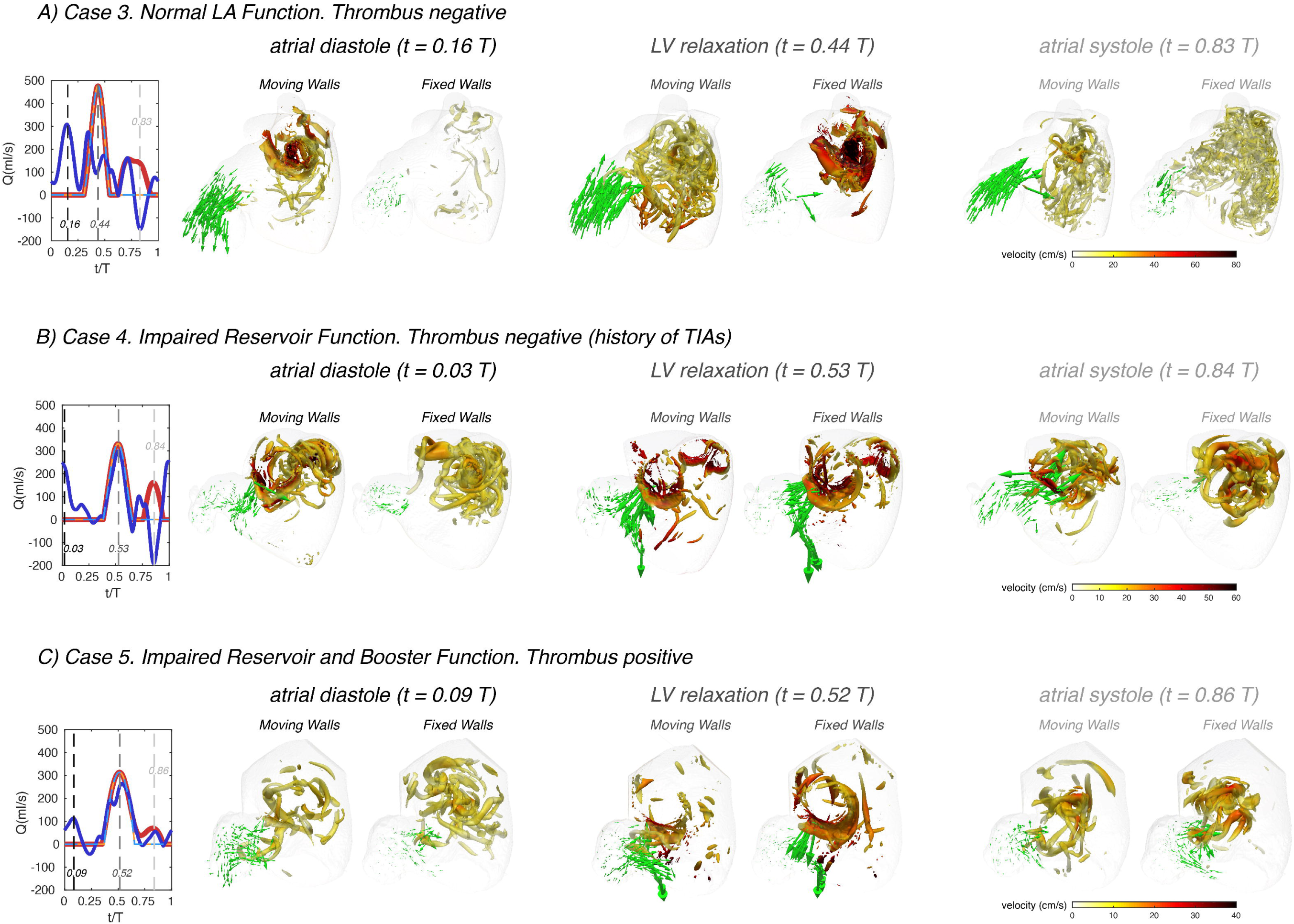
Left atrial vortex patterns and left atrial appendage flow velocity in subjects with normal and impaired atrial function. Three subjects are shown, one with normal atrial function (**panel A**, same subject as in Figure 2), one with impaired reservoir function (**panel B**), and one with both impaired reservoir and booster functions (**panel C**). For each subject, instantaneous 3D flow visualizations are shown in the frontal view of the LA using data the moving-walls and fixed-walls simulations. In each visualization, we rendered the flow velocity vectors in the LAA vicinity and vortex cores in the whole LA (as detected by thresholding the second invariant of the velocity gradient tensor, *Q = 1000* s^-2^). The LAA velocity vectors were amplified by a factor of 4 with respect to those in Figure 2, and the surface of the vortex cores was colored according to the local absolute value of blood velocity, as indicated in the colorbars. Each panel includes plots of the time histories of flow rate through the mitral valve (red) and the cumulative flow rate through the pulmonary veins (blue), both for the moving-walls (dark color, thick continuous lines) and fixed-walls (light color, thin dashed lines) simulations. For each subject, flow visualizations are shown at three instants of time. The first one corresponds to atrial diastole (i.e. peak flow rate through the pulmonary veins in the moving wall simulations, *t/T = 0.16, 0.03* and *0.09* in panels A, B and C). The second instant of time corresponds to left ventricular rapid filling (i.e. the E-wave, peak flow rate through the mitral valve, *t/T = 0.44, 0.53* and *0.52* in panels A, B and C). The third instant of time corresponds to atrial systole (i.e. the A-wave of left ventricular filling and peak backflow rate through the pulmonary veins, *t/T = 0.83, 0.84* and *0.86* in panels A, B and C). These three instants of time are indicated with black, dark grey and light grey dashed vertical lines in the flow-rate plots at the left-hand-side of each panel.

The subject with normal atrial function exhibits two vortex-ring systems associated with the atrial filling jets discharging from the left and right PVs (Figure 3A). These vortex rings collide into each other and break down into a complex network of fine-scale vortices that fills the whole atrial body, and which is sustained throughout the cardiac cycle. The fixed-wall simulation presents two strong vortex-ring systems that break down into finer-scale vortices, but these decay almost completely by atrial diastole of the next cardiac cycle. As shown in Figure 2, the magnitude and dynamics of the velocity vectors in the LAA of this normal subject vary markedly between the moving-wall and fixed-wall simulations.

In patients with impaired atrial function, moving-wall and fixed-wall simulations yield qualitatively similar LA vortex systems, while the velocity fields inside the LAA exhibit interesting differences. In patient 4 (Figure 3B), who had severely impaired reservoir function (passive LA ejection fraction = 0.04, see Table 1), the LA wall was akinetic during atrial diastole and LV relaxation. Consequently, the moving-wall and fixed-wall simulations produced similar LAA velocity patterns during these phases. However, this patient had a relatively normal booster function (booster LA ejection fraction = 0.2) with normal LA wall kinetics during atrial systole. As a result, blood velocity in the LAA showed and outward flow pattern for the moving-wall simulations whereas it showed a weak recirculating motion for the fixed-wall simulations. In subjects with decreased reservoir and booster function, *e.g.*, subject 5 in Figure 3C, the moving-wall and fixed-wall simulations produced qualitatively similar LAA velocity distributions across the whole cardiac cycle. Consistent with patient 5 having more passive than booster LA function, the most appreciable difference between fixed-wall and moving-wall simulations was observed for atrial diastole – during this phase, the moving-wall LAA has a weak inward-flowing pattern in addition to the swirl that is found in the fixed-wall LAA.

### 3.4. The effect of atrial function in residence time and kinetic energy

To visualize the effect of impaired atrial function on blood stasis inside the LA chamber, we mapped the blood residence time in two oblique intersecting plane sections of the LA body and the LAA. These maps are represented in Figure 4 for the same three study subjects of Figure 3, both for moving and fixed walls. Once converged, the *T*_*R*_ maps become quasi-periodic, and show that blood is systematically more stagnant inside the LAA than in the LA body. Residence time gradients are also significantly sharper inside the LAA, consistent with flow stirring being lower in the appendage than in the atrial body.

**Figure 4.**
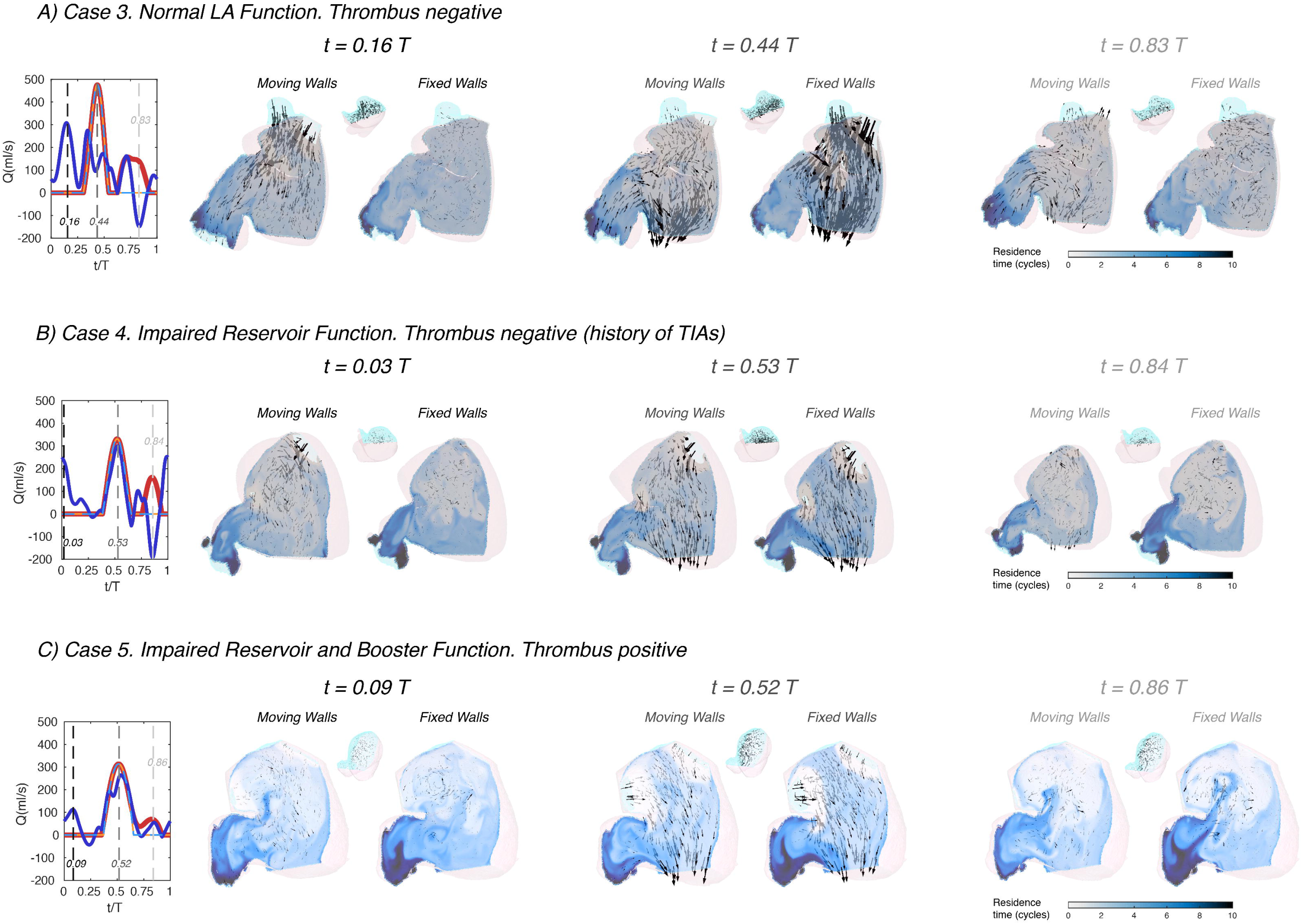
Left atrial blood residence time in subjects with normal and impaired atrial function. Three subjects are shown, one with normal atrial function (**panel A**), one with impaired reservoir function (**panel B**), and one with both impaired reservoir and booster functions (**panel C**). Each subject includes data from both moving-walls simulations and fixed-walls simulations. For each subject, instantaneous 2D fields of the absolute value of the residence time *T*_*R*_ (see main text for definition) are plotted in two oblique plane sections of the LAA and the LA body that separate the cyan and pink parts of the sagittal view insets. Additionally, 3D blood velocity vectors are rendered behind the sectioning planes (i.e. inside the cyan part of the LAA and the LA body). Similar to Figure 3, each panel includes plots of the time histories of flow rate through the mitral valve (red) and the cumulative flow rate through the pulmonary veins (blue), both for the moving-walls (dark color, thick continuous lines) and fixed-walls (light color, thin dashed lines) simulations. The data are represented at three instants of time. The first one corresponds to atrial diastole (i.e. peak flow rate through the pulmonary veins in the moving wall simulations, *t/T = 0.16, 0.03* and *0.09* in panels A, B and C). The second instant of time corresponds to left ventricular rapid filling (i.e. the E-wave, peak flow rate through the mitral valve, *t/T = 0.44, 0.53* and *0.52* in panels A, B and C). The third instant of time corresponds to atrial systole (i.e. the A-wave of left ventricular filling and peak backflow rate through the pulmonary veins, *t/T = 0.83, 0.84* and *0.86* in panels A, B and C). These three instants of time are indicated with black, dark grey and light grey dashed vertical lines in the flow-rate plots at the left-hand-side of each panel.

Visual inspection of the *T*_*R*_ maps suggests that LAA blood stasis in LAAT/TIA-neg patients (Figure 4A) was less prominent than in LAAT/TIA-pos patients (Figure 4B-C). To confirm this result, we compiled the statistical distributions of LAA *T*_*R*_ for all of our subjects using 200 time instants equally distributed over the cardiac cycle. We also compiled statistics of the kinetic energy *K = 1/2 (u*^*2*^ *+ v*^*2*^ *+ w*^*2*^), since this hemodynamic variable is more accessible via medical imaging than *T*_*R*_. The data from moving-wall simulations show that LAA residence time increases as *EF*_*LA*_ worsens across different subjects (Figure 5A). Although less clear, we also found a trend for K to decrease as *EF*_*LA*_ worsens (Figure 5B). In the fixed-wall simulations all subjects have zero ejection fraction, but the data suggests that subjects with higher LA volume tend to have higher *T*_*R*_, and lower *K* (Figure 5C-D).

**Figure 5.**
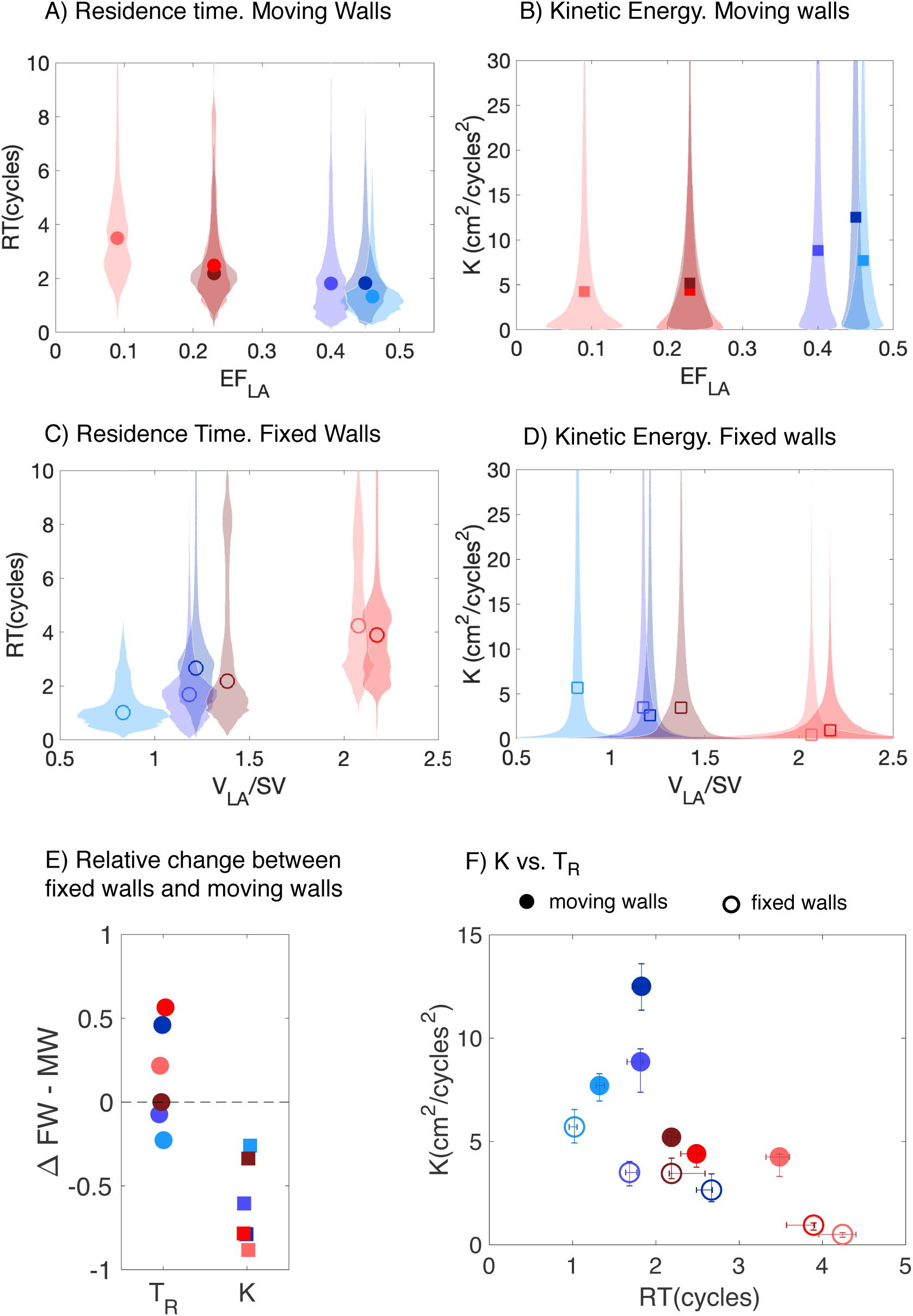
Probability distributions of residence time and instantaneous kinetic energy inside the left atrial appendage of subjects with normal and impaired atrial function. **A, B)** Violin plots of the probability density function (shaded patches) of *T*_*R*_, and *K* inside the LAA of the N = 6 subjects studied, together with their median (symbols). The data come from the moving-wall simulations and are plotted vs. the LA ejection fraction of each subject. **C, D)** Violin plots of the probability density function of the same variables as obtained from fixed-wall simulations. The data are plotted vs. the volume of the LA normalized by stroke volume. **E)** Relative change in mean values of *T*_*R*_,and *K* between the fixed-wall (FW) and the moving-wall (MW) simulations, defined as 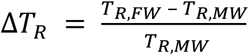, etc. **F)** Scatter plot of *K* vs *T*_*R*_ for moving-wall (solid symbols) and fixed-wall (open symbols) simulations. The error bars in each panel span the 95% confidence intervals of the mean. Solid and hollow symbols come respectively from moving-wall and fixed-wall. In all the panels, each subject’s data is colored using the same scheme used in Figure 1 (blue shades indicate LAAT/TIA-neg patients and red shades indicate LAAT/TIA-pos patients).

### 3.5. Hemodynamic parameters and thrombosis risk in moving-wall and fixed-wall simulations

Figure 5A-B demonstrates that blood inside the left atrial appendage of LAAT/TIA-pos patients had marked alterations in hemodynamic parameters when compared with LAAT/TIA-neg patients. Specifically, LAAT/TIA-pos patients had higher residence time and lower kinetic energy inside their LAA than LAAT/TIA-neg patients. Given that these trends seemed to hold at least in part for the fixed-wall simulations (Figure 5C-D), we examined the variation in LAA *T*_*R*_ and *K* between fixed-wall and moving-wall simulations (Figure 5E). This analysis showed that *K* decreased in the fixed-wall simulations across all study subjects; however, the variations in *T*_*R*_ were not systematic. In particular, fixed-wall simulations yielded equal or lower LAA *T*_*R*_ than moving-wall simulations for two of the three LAAT/TIA-neg subjects. While these results are not easy to reconcile with the dependence of *T*_*R*_ on *EF*_*LA*_ (Figure 5A), they are consistent with our finding that fixed-wall simulations tend to represent the LA and LAA flow patterns better in atria with impaired function than in normal atria (Figure 2). Finally, we examined whether fixed-wall simulations were less accurate in separating the LAAT/TIA-pos and LAAT/TIA-neg groups in our small study population. Scatter plots of LAA *T*_*R*_ and *K* (Figure 5F) showed some overlap between the LAAT/TIA-pos and LAAT/TIA-neg groups in the fixed-wall simulations. In contrast, moving-wall simulations provided good clustering of the groups for either *T*_*R*_ or *K*.

## 4. Discussion

Thrombogenesis is most likely to occur with the concurrence of endothelial damage or dysfunction, blood stasis, and the presence of procoagulatory factors in blood. Blood flow governs the transport of the enzymes, chemical factors, and platelets to and from endothelial damage sites [6]. Blood stasis permits the reactive species to remain close enough to each other to initiate clot formation [25]. The goal of this study was to investigate how LA function influences LAA blood stasis. More specifically, we hypothesized that CFD analyses of LA/LAA hemodynamics could identify patients at increased risk of LAA thrombosis.

### 4.1. Atrial Function, Flow Patterns, and Blood Stasis

By selecting subjects with a wide range of LA ejection fractions (*EF*_*LA*_), we characterized how LA and LAA flow patterns vary with atrial function. Consistent with previous MRI and CFD reports of flow in the LA [7, 14, 28], we found that blood from the PVs flows directly towards the mitral annulus during ventricular diastole, whereas a large recirculating pattern forms inside the LA body during ventricular systole. In subjects with impaired reservoir function, reduced wall motion during ventricular systole caused slower recirculation and quicker dissipation of fine-scale vortices.

In the LAA, normal wall kinetics drove inward and outward flow jets. By contrast, flow inside the LAA was similar to a lid-driven cavity flow in subjects with impaired booster and reservoir function, consistent with Masci *et al*’s [19] data. Additionally, our subject with impaired reservoir function and normal booster function experienced hybrid LAA flow dynamics that oscillated between lid-driven cavity flow during LA diastole, and a weak outflow jet during LA systole. Taken together, our results indicate that flow patterns inside the LAA are not as complex as in the LA body or the LV, which we attribute to the strong geometrical confinement of the flow inside the appendage. These flow patterns led to elevated blood residence in the LAA compared to the LA body, in agreement with previous CFD results [3, 8, 15, 18, 19, 22], and the consensus that thrombi form preferentially in the LAA. Patients with impaired atrial function also showed a trend for blood stasis to appear near the mitral annulus, since their weaker LA body vortex systems caused decreased stirring compared to normals.

### 4.2. Blood Transit and LAA Thrombogenesis

While previous CFD studies suggested that reduced wall motion impairs LA blood transit [18, 19, 22], there is a lack of systematic data about how functional parameters such as *EF*_*LA*_ affect this transit. We demonstrated that LAA *T*_*R*_ has a clear dependence on *EF*_*LA*_. We also showed how LAA *T*_*R*_ is related to the risk of thrombosis, leveraging that our database includes two subjects with LAA thrombi and one with a history of TIAs. We and others have recently shown that LV blood stasis in the setting of myocardial infarction precedes LV mural thrombosis [9, 17] and cerebral microembolism [4]. The present study extends this idea to the LAA showing that LAA thrombosis could occur when the mean residence time inside the LAA exceeds ∼2 s (*i.e.*, 2 cycles). This threshold agrees with Martinez-Legazpi et al’s [17] threshold for LV thrombosis, which is notable considering the different etiologies of myocardial infarction and AF, and the different methodologies used in our two studies.

In addition to residence time, we show that kinetic energy could be potentially useful to predict the risk of LAA thrombosis. This knowledge could be useful because, while *T*_*R*_ is a cumulative variable whose calculation requires solving a partial differential equation, calculating *K* is significantly less involved. In particular, *K* should be accessible by medical imaging modalities with moderate time resolution, such as phase-contrast MRI. Nevertheless, the observed predictive ability of *K* in the case of the LAA could be in large part due to the unique geometrical characteristics of the appendage and its particular location within the LA. We note that *K* can yield poor predictions of blood stasis in other scenarios [11].

### 4.3. Fixed-Wall vs. Moving-Wall Simulations to Predict Thrombogenesis Risk

Fixed-wall simulations provided flow fields and stasis maps that differed from those obtained in moving-wall simulations, especially in subjects with normal atrial function. This discrepancy is not surprising; in addition to neglecting LA wall motion, fixed-wall simulations have oversimplified inflow-outflow boundary conditions and ignore LV – LA interactions. Despite these limitations, we found that fixed-wall simulations could still be useful to assess the risk of thrombogenesis on a patient-specific basis. This knowledge may be of value because fixed-wall simulations only require static images which are more commonly available in clinical practice, and are less involved from the point of image segmentation and CFD analysis. In our reduced study cohort, *T*_*R*_ and *K* in the fixed-wall LAA were only slightly less accurate predictors of LAA thrombus or TIAs than their moving-wall counterparts.

While these findings warrant further evaluation in larger patient cohorts, they reflect the influence of LA and LAA geometry on blood flow. Morphological analysis of the LA chamber in our subject cohort (Table 1) revealed that the LAAT/TIA-pos subjects had the largest LA volumes and showed a trend for higher LA sphericity, consistent with clinical reports of stroke rates vs. LA/LAA geometry in AF patients [2, 5, 29]. Moreover, our LAAT/TIA-pos subjects, which had the highest LAA residence time in both our moving-wall and fixed-wall simulations, also had LAA morphologies that are ranked more pro-thrombogenic according to DiBiase *et al*’s categorization [5]. Although the number of LAA lobes measured by 3D ultrasound imaging and stroke risk have been shown to correlate AF patients [29], we did not observe a relationship between LAA lobe number and *T*_*R*_, K or presence of LAAT/TIAs (Table 2). This discrepancy could be caused by our small cohort size and differences in spatial resolution between CT and ultrasound.

### 4.4. Strengths and Limitations of the Present Study

The number of patients considered in this study was small (N = 6), even considering that we performed simulations with both moving and fixed walls for each subject. However, our patient cohort was diverse enough to explore how different aspects of LA function (e.g., reservoir, booster pump, and conduit) affect the flow inside this chamber with an unprecedented level of detail. Also, while small, the number of subjects considered here is higher than in most previous CFD studies [3, 18, 22]. More importantly, by including patients without LAA thrombus and with LAA thrombus or history of TIAs, we were able to demonstrate that CFD analysis of LA flow could predict the risk of LAA thrombogenesis.

We did not consider a model of the coagulation cascade in this study. Biochemical modeling of thrombogenesis could be incorporated into the CFD framework [25]. However, the additional models would markedly increase the cost of the simulations. Also, the accuracy of these models is unclear, given their dependence on many reaction constants whose values are difficult to obtain in a patient-specific basis.

We considered the same heart rate and blood viscosity for all patients to eliminate these parameters as independent variables in our analysis, even if these parameters may vary from patient to patient. In LAAT-pos patients, we segmented the thrombus and removed it before performing the CFD analyses in an effort to recapitulate the conditions that caused thrombosis. However, the presence of a thrombus could constrain the motion of the LAA walls – thus, pre-thrombus LAA wall motion might have not been fully recovered. One of our subjects had a history of TIAs of unknown origin, which we presumed to be cardioembolic from the LA/LAA, but we cannot exclude the possibility of another etiology.

Similar to previous simulations [3, 8, 15, 18, 22], our LA model did not consider the mitral valve in the outflow of the chamber. This choice is justified by whole left heart CFD studies showing that the mitral valve does not significantly affect LA flow [28]. Our work improves over most previous computational studies of the LA flow [3, 8, 18] by considering patient-specific inflow/outflow boundary conditions, similar to Otani *et al* [22]. One limitation of our simulations is that PV flow rates were evenly distributed [16]. More patient-specific modeling of inflow boundary conditions would involve measuring the flow rate through each pulmonary vein (e.g., by transesophageal echocardiography) or modeling the flow impedance of each pulmonary vein based on imaging data such as vein diameter [26].

As discussed above, fixed-wall simulations have severe limitations as a model of AF, including but not limited to their inability to capture LA motion and deformation mediated by the chamber’s passive response to incoming PV flow and LV-LA interactions. The rationale for including fixed-wall simulations in our study was to assess how these limitations affect the prediction of patient-specific hemodynamic parameters such as residence time, by comparing with moving-wall simulations. This assessment is warranted considering that several previous studies used fixed-wall simulations [3, 8].

In conclusion, this work combined 4D-CT acquisitions of left atrial wall motion with immersed-boundary CFD analyses of LA blood flow to establish proof of principle of personalized risk stratification of LAA thrombogenesis for patients with AF that could be translated to the clinical setting. For the large number of AF patients for whom there is currently uncertainty regarding optimum medical treatment, the proposed risk stratification paradigm may allow clinicians to better balance the benefits of chronic anticoagulation therapy (or LAA occlusion therapy) with the hemorrhagic risks of therapy. Larger studies are warranted to evaluate the robustness and potential clinical utility of this methodology.

## Notes

### Competing Interest Statement

The authors have declared no competing interest.

